# Heterotypic FG-Nup98-tau condensates form nested assemblies through stoichiometry-dependent phase transitions

**DOI:** 10.64898/2026.05.22.727239

**Authors:** Niharika Nag, Sumangal Roychowdhury, Amar Jeet Yadav, Aditya Kumar Padhi, Krishnananda Chattopadhyay, Timir Tripathi

## Abstract

The dysfunction of nucleocytoplasmic transport (NCT) and tau aggregation are emerging as interconnected hallmarks of neurodegenerative tauopathies. However, the molecular basis by which components of the nuclear pore complex and tau interact remains unclear. Here, we combine experimental and computational approaches to elucidate the mechanism of heterotypic phase separation between the FG-repeat domain of nucleoporin Nup98 (FG-Nup98) and tau. *In vitro* assays revealed that FG-Nup98 and tau undergo coacervation, forming dynamic condensates whose morphology and dynamics depend on stoichiometry, macromolecular crowding, and ionic strength. FRAP indicated reduced tau mobility within FG-Nup98-rich condensates, supporting a scaffold-client model. Complementary computational analyses revealed hierarchical binding energetics: FG-Nup98 self-association is strongest, followed by FG-Nup98-tau and tau-tau interactions. While FG-Nup98 forms stable homotypic networks, tau-tau contacts are transient but energetically favorable, which suggests that elevated tau concentrations may trigger a transition from droplets to tau aggregates. Together, these results establish that multivalent FG-Nup98-tau interactions drive condensate formation that could potentially perturb the permeability barrier of the nuclear pore. This study elucidates the coordinated behaviors of FG-Nup98-tau condensates and provides a framework for understanding NCT defects in tauopathies.

## Introduction

Neurofibrillary tangles, composed of aggregated tau protein, are a defining neuropathological hallmark of Alzheimer’s disease (AD) and related tauopathies. Tau, a microtubule-associated protein, becomes hyperphosphorylated in disease states, leading to its pathological self-assembly into insoluble fibrils within neurons ^1^. The progression of tau aggregation across brain regions closely correlates with synaptic dysfunction and neurodegeneration, implicating tau as a central driver of cognitive decline ^2, 3^. Although the structural features of tau aggregates have been extensively studied, the cellular triggers and molecular mechanisms that initiate tau misfolding and assembly remain incompletely understood.

Recent findings suggest an unexpected link between tau pathology and the nuclear pore complex (NPC), the essential gateway for nucleocytoplasmic transport (NCT) ^4, 5^. In AD and other tauopathies, nucleoporins, including the FG-repeat nucleoporin Nup98, are mislocalized from the nuclear envelope to the cytoplasm, where they co-aggregate with tau ^4, 6, 7^, causing morphological distortion of the NPCs and a disruption in the NCT, thus leading to neurodegeneration. *In vitro* studies further indicate that the C-terminal FG-repeat domain of Nup98 directly accelerates tau fibrillization, suggesting a synergistic role in tau aggregation ^4, 8^. Tau and Nup98 share key biophysical features: both contain intrinsically disordered, low-complexity domains that mediate weak, multivalent interactions and are each individually capable of undergoing liquid-liquid phase separation (LLPS) ^9, 10, 11^. The microtubule-binding and proline-rich regions of tau promote droplet formation *in vitro*, which may act as precursors to fibril formation ^12^. Similarly, the FG-repeat domain of Nup98 forms phase-separated condensates and hydrogel-like barriers within the NPC ^13, 14^. Recent surface plasmon resonance studies have confirmed that tau interacts directly and with high affinity with FG-repeat domains of Nup98, primarily via the positively charged repeat region of tau ^8^. All these observations suggest that tau and Nup98 may co-assemble into heterotypic condensates that concentrate tau and promote its pathological conversion.

Here, we integrate experimental and computational approaches to investigate the interaction between tau and the FG repeat domain of Nup98. Using *in vitro* droplet assays, confocal microscopy, and fluorescence recovery after photobleaching (FRAP), we characterized the phase behavior of the condensates formed by tau and FG-Nup98. Molecular docking and multiscale molecular dynamics simulations further identify key interaction sites and the energetic features of this disordered interface. Together, our findings support a mechanistic model in which aberrant phase separation between tau and FG-Nup98 may facilitate tau aggregation and eventually contribute to nucleocytoplasmic transport defects in tauopathy. The pharmacological targeting of the tau and Nup98 interaction is a strategy being explored to restore nucleocytoplasmic transport and nuclear pore complex integrity in AD and related tauopathies ^5^. The molecular basis of this interaction provides insight into a potential driver of neurodegeneration and identifies a targetable axis in disease progression.

## Results

### Bioinformatic analysis reveals intrinsic disorder and phase separation potential of FG-Nup98 and tau

We first asked whether the primary sequence features of the FG-repeat domain of Nup98 and tau are intrinsically suited for spontaneous phase separation. To assess this, we performed a comprehensive bioinformatic analysis integrating multiple disorder and phase separation predictors. Using the RIDAO ^15^ meta-server, we found that both proteins are almost entirely intrinsically disordered. Notably, tau displayed particularly high disorder in its N-terminal and repeat-rich regions, consistent with previous reports (Supplementary Figure S1A). FG-Nup98, comprising the phenylalanine-glycine repeat domain of Nup98, was also predicted to be disordered across its full length (Supplementary Figure S1B). We next analyzed the electrostatic properties of the two proteins using the CIDER platform (https://pappulab.wustl.edu/CIDER/) ^16^, which showed that tau exhibited a strongly positive NCPR in its microtubule-binding repeat region and FG-Nup98 displayed an overall mildly positive NCPR (Supplementary Figure S1C and S1D). To further probe their capacity for phase separation, we applied the FuzDrop server ^17^ to compute their LLPS propensities. FG-Nup98 and tau received pLLPS scores of 1.000 and 0.9951, respectively, which indicated extremely high intrinsic likelihood to undergo liquid-liquid phase separation (Supplementary Figure S1E and S1F). These findings collectively suggested that FG-Nup98 and tau harbor sequence-level features that could be conducive to undergoing co-condensation.

### Stoichiometry governs the condensate architecture of FG□Nup98-tau

To convert our bioinformatic predictions into a droplet formation profile, we screened FG-Nup98 and tau across a concentration matrix (0-20□µM each) under crowded conditions. Turbidity measurements at 340□nm showed that, even without polyethylene glycol (PEG), wells containing ≥□2□µM FG-Nup98 turned turbid, whereas tau alone remained largely soluble; adding 5-10□% PEG lowered the tau threshold and amplified the turbidity across the entire matrix, revealing a concentration and crowding-dependent phase space (Figure 1A). Confocal microscopy offered a closer look at the condensate architecture generated under these conditions, wherein two mutually exclusive morphologies emerged. The LLPS of the individual proteins were verified (Supplementary Figure S1G and S1H) before moving on to the protein mixtures. When tau was present in large excess (≥□10:1), we observed uniformly mixed, colocalized droplets in which tau (green) and FG-Nup98 (red) occupied a single dense phase. As the relative abundance of FG-Nup98 increased, the system reorganized into “nested” condensates, where red FG-Nup98 sub-droplets sat within larger green tau droplets (Figure□1B). The transition between the architectures seemed to be governed by the stoichiometry of the two proteins. On constructing a concentration-ratio phase diagram (Figure□2), we found that no condensates formed when both proteins were at 2□µM, defining a minimal nucleation threshold. Above this limit, ratios of ≥□10:1 produced exclusively colocalized droplets; ratios between 5:1 and 2:1 yielded a mixed population in which tau-rich colocalized condensates coexisted with nested structures; and ratios at or below 1:1 favoured either nested condensates or isolated FG-Nup98 droplets. Pushing FG-Nup98 beyond 10□µM ultimately drove the red droplets to coarsen into irregular aggregates, suggesting that an excess of FG-Nup98 leads to a clustering of the droplets and the formation of aggregate-like structures. Taken together, these data demonstrate that macromolecular crowding lowers the energetic barrier for FG-Nup98-tau phase separation, while the relative stoichiometry of the two proteins dictates whether they share a common dense phase or demix into separate droplets-within-droplets. Such a tunable organisation parallels the heterogeneous droplet architectures reported for other neurodegeneration-linked proteins ^18^ and sets the stage for dissecting how composition influences material properties and downstream aggregation.

**Figure 1:**
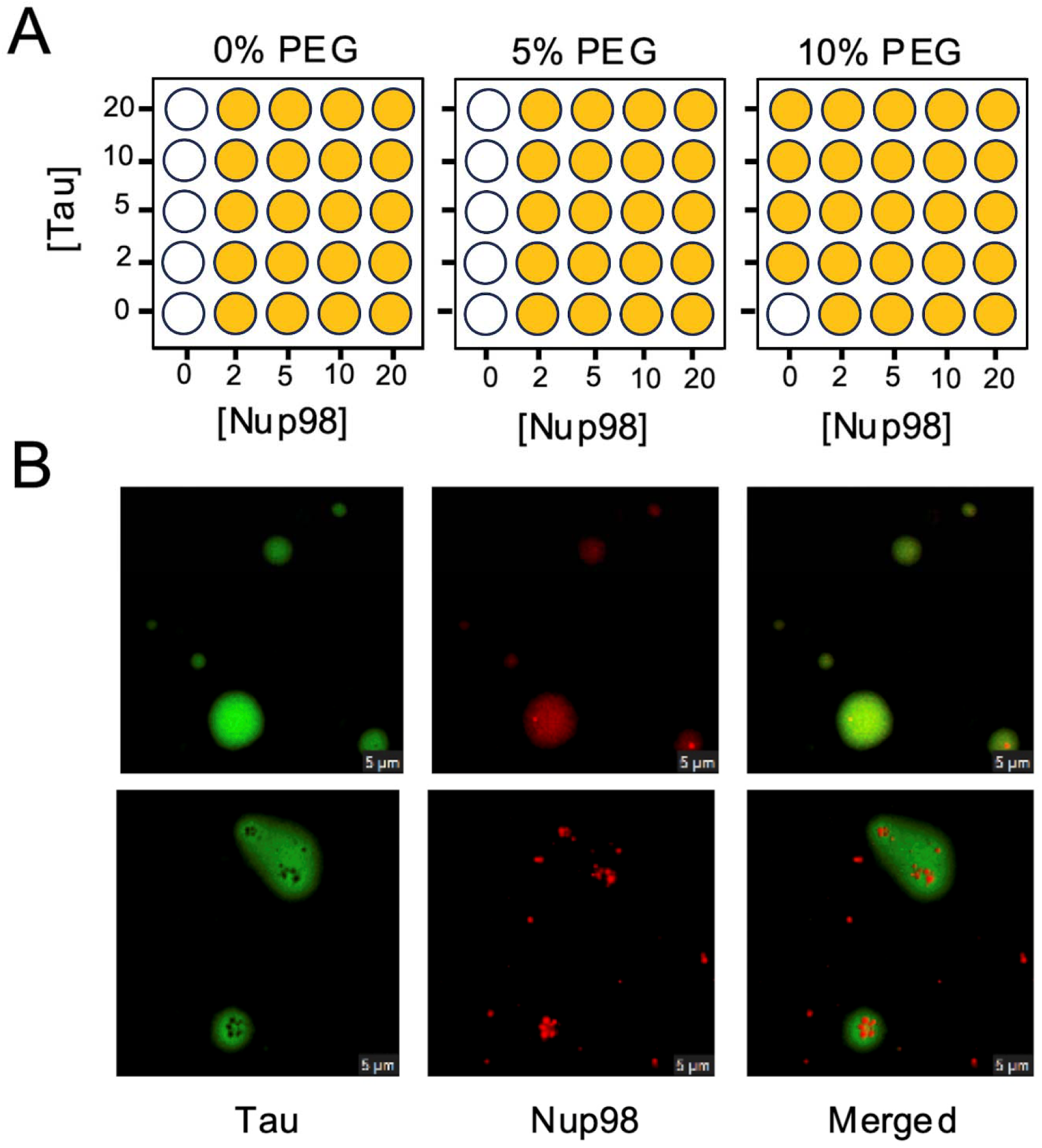
Phase separation of FG-Nup98 and tau. (A) Turbidity assay at OD340 showing phase separation at varying FG-Nup98 and tau concentrations with 0%, 5%, and 10% PEG. Filled circles indicate turbidity >0.15, depicting droplet formation. (B) Confocal images showing two types of FG-Nup98-tau droplets. Top row: colocalized droplets where both proteins mix. Bottom row: nested droplets where FG-Nup98 forms distinct condensates within tau droplets.

**Figure 2:**
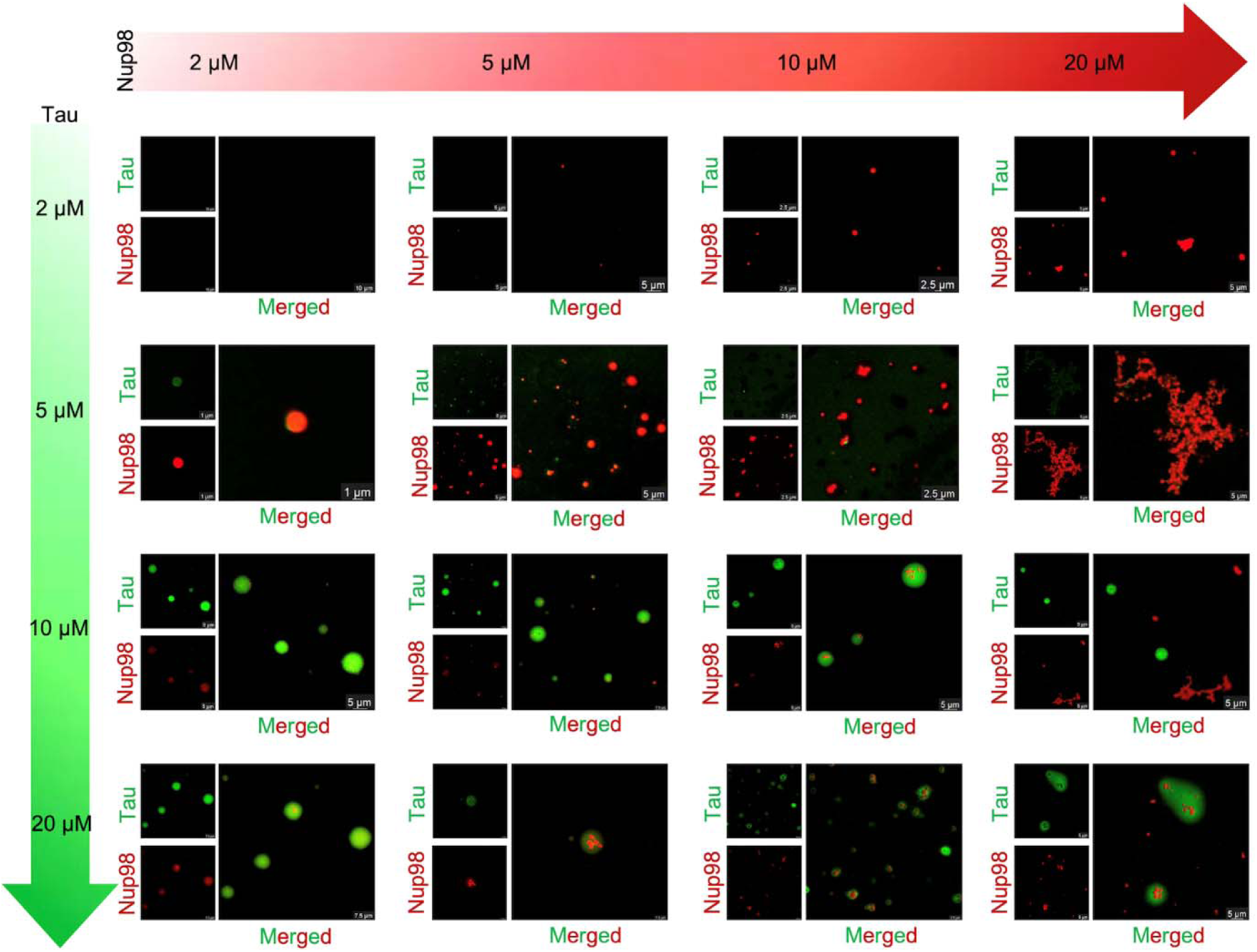
Phase diagram of FG-Nup98 and tau co-condensation. Confocal microscopy images showing tau (green) and FG-Nup98 (red) phase separation at different protein concentrations. The x-axis represents increasing FG-Nup98 concentration (2-20 μM), while the y-axis represents increasing tau concentration (2-20 μM).

### PEG and ionic strength remodel FG□Nup98-Tau condensates

We fixed the protein stoichiometry at 10:1 (20□µM tau, 2□µM FG□Nup98) and then varied the concentrations of crowding agent and salt to see how the physical environment affected the condensates. Without PEG, no phase separation was observed, but 5□% PEG produced a heterogeneous field in which most droplets were colocalized while a minority displayed small, FG□Nup98-rich inclusions inside tau□rich hosts. Raising the concentration of crowder to 10□% homogenised the system, and nearly every droplet became uniformly mixed and spherical (Figure 3A). Keeping the PEG concentration at 10□%, we then varied the concentration of NaCl. In the presence of 0□mM salt, the droplets were fully co-localized and liquid-like. Introducing 150□mM NaCl, roughly physiological strength, yielded a mixed population: intact green□and□red droplets coexisted with separate FG□Nup98 and tau droplets, and many tau□rich condensates adopted irregular shapes. In the presence of 500□mM NaCl, FG□Nup98 and tau fused into amorphous clusters, and at 1□M NaCl, only red FG□Nup98 aggregates persisted, indicating that high ionic strength disrupts the heterotypic interactions that keep tau in the dense phase (Figure□3B). This trend is consistent with the intrinsic salt sensitivities of the two proteins: tau’s liquid□liquid phase separation is promoted at low ionic strength and suppressed as salt rises, whereas FG□ Nup98 phase separation is actually enhanced by higher salt concentrations ^19, 20, 21, 22^. The data suggest that modest crowding plus low□to□moderate salt stabilise a shared, liquid condensate. Aggressive charge screening seemed to push towards demixing and eventual aggregation. Such environmental tunability suggests that subtle shifts in the cellular milieu could tip FG□Nup98-tau assemblies from functional droplets to pathogenic solids.

**Figure 3:**
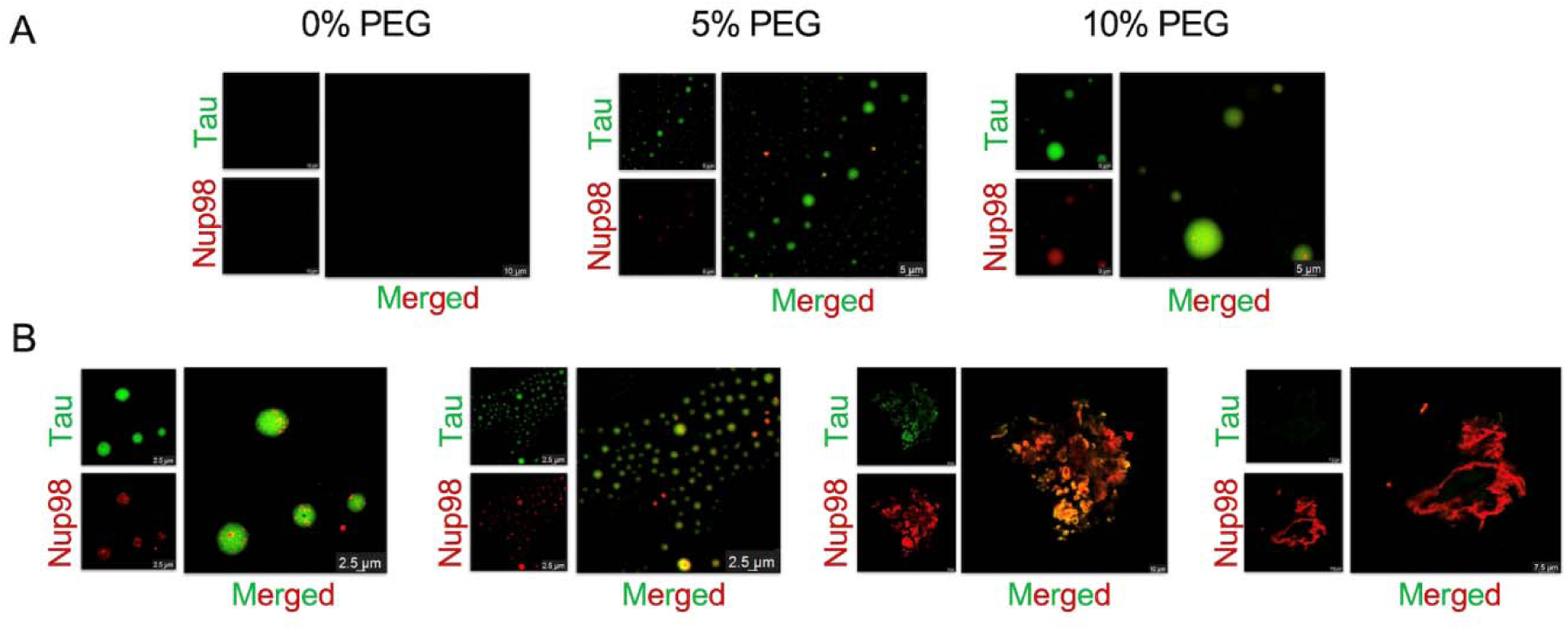
Effect of PEG and salt on FG-Nup98-tau phase separation. (A) Confocal microscopy images showing tau (green) and FG-Nup98 (red) droplets at varying PEG concentrations (0%, 5%, 10%) and 50 mM NaCl. (B) FG-Nup98-tau droplets at 10% PEG with increasing NaCl concentrations (0 mM, 150 mM, 500 mM and 1 M) showing that higher salt leads to aggregation and altered droplet structures.

**Figure 4:**
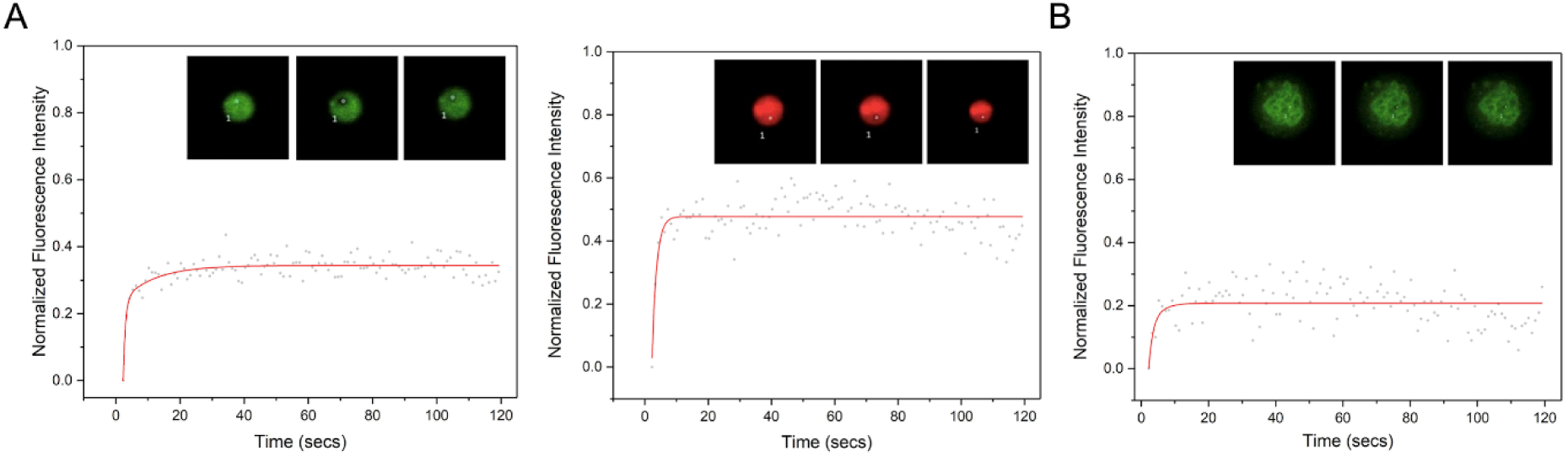
Fluorescence Recovery After Photobleaching (FRAP) analysis of FG-Nup98 and tau droplets. (A) FRAP analysis of colocalized tau (green) and FG-Nup98 (red) droplets. (B) FRAP analysis of tau in nested droplets.

### FG-Nup98 binding redirects tau mobility and dynamics

It has been reported in a previous study by Lisa Diez et al. that tau binds to FG-Nup98 with a dissociation constant (K_D_) of 0.4 μM, indicating that the two disordered chains can form substantial heterotypic contacts even outside a condensed phase ^8^. To ask how this interaction reshapes the diffusional dynamics of the droplets, we photobleached labelled proteins in both colocalized and nested condensates. In mixed, colocalized droplets, tau recovered to ∼34□,% of its pre-bleach intensity, while FG-Nup98 recovered to ∼48□,%, confirming that both partners remain largely mobile (Figure□,4A, Supplementary Video 1 and 2). In nested droplets, however, tau recovery fell to ∼21□,% and the FG-Nup98 core could not be measurably bleached, implying that the inner FG-Nup98 compartment acts as a rigid scaffold that restricts tau diffusion (Figure□,4B, Supplementary Video 3).

### Assessment of binding affinities and energetics from protein-protein docking experiments

To complement our biophysical data with atomistic detail, we first generated monomeric models of the FG repeat domain of Nup98 and tau using AlphaFold□v2.0. Only the top□ranked structures, each displaying high overall confidence and pLDDT scores, were utilized for molecular docking (Supplementary Figure S2). We analyzed the docked conformations, binding affinities, and interaction energetics of the FG-Nup98-FG-Nup98, FG-Nup98-tau, and tau-tau complexes to assess their stability and binding strength. These analyses revealed that the FG-Nup98-FG-Nup98 complex had the highest binding affinity, with GOAP, DFIRE, and ITScore values of –25270.73, –25724.44, and –12118.18 kcal/mol, respectively, and a Ranksum of 17. In comparison, the FG-Nup98-tau complex showed moderate affinity, with GOAP, DFIRE, and ITScore scores of –18989.09, –19681.47, and –9188.73 kcal/mol, and a Ranksum of 33. The tau-tau complex demonstrated the lowest affinity, recording GOAP, DFIRE, and ITScore values of – 12711.30, –13690.87, and –6086.16 kcal/mol, along with a Ranksum of 19 (Supplementary Table S1).

These results rationalise our microscopy data; the strong FG-Nup98-FG-Nup98 affinity promotes the formation of dense FG□,Nup98 cores, which can act as scaffolds around which tau concentrates. Even in colocalized droplets, FG□,Nup98 frequently concentrated in regions, suggesting that tau-tau interactions are weaker and insufficient to dominate condensate organization in the presence of Nup98. A detailed structural analysis of the FG-Nup98-FG-Nup98 complex identified key interacting residues. Specifically, N95 forms polar contacts with A456, Q76 forms polar contact with P457, while M1 and G2 establish two hydrogen bonds with A348, further stabilizing the FG-Nup98-FG-Nup98 complex (Figure 5A-C). In the FG-Nup98-tau complex, key intermolecular interactions were identified, which contribute to the complex’s stability. S250 of FG-Nup98 forms a polar contact with K140 of tau, while T166 of FG-Nup98 establishes two hydrogen bonds with K130 and K132 of tau. Additionally, Q193 and S185 of FG-Nup98 interact via a polar contact with V128 and E117 of tau, respectively (Figure 5D-F). In contrast, the tau-tau complex displayed the fewest hydrogen-bond interactions. Notably, E338 formed two hydrogen bonds with T319, and G60 engaged in a polar contact with D81(Figure 5G-H). The computational results mirror our experimental findings that FG□,Nup98 is predisposed to self□,assemble, while its interaction with tau, though weaker, is still strong enough to drive heterotypic condensation, reinforcing the reliability of the docking model.

**Figure 5:**
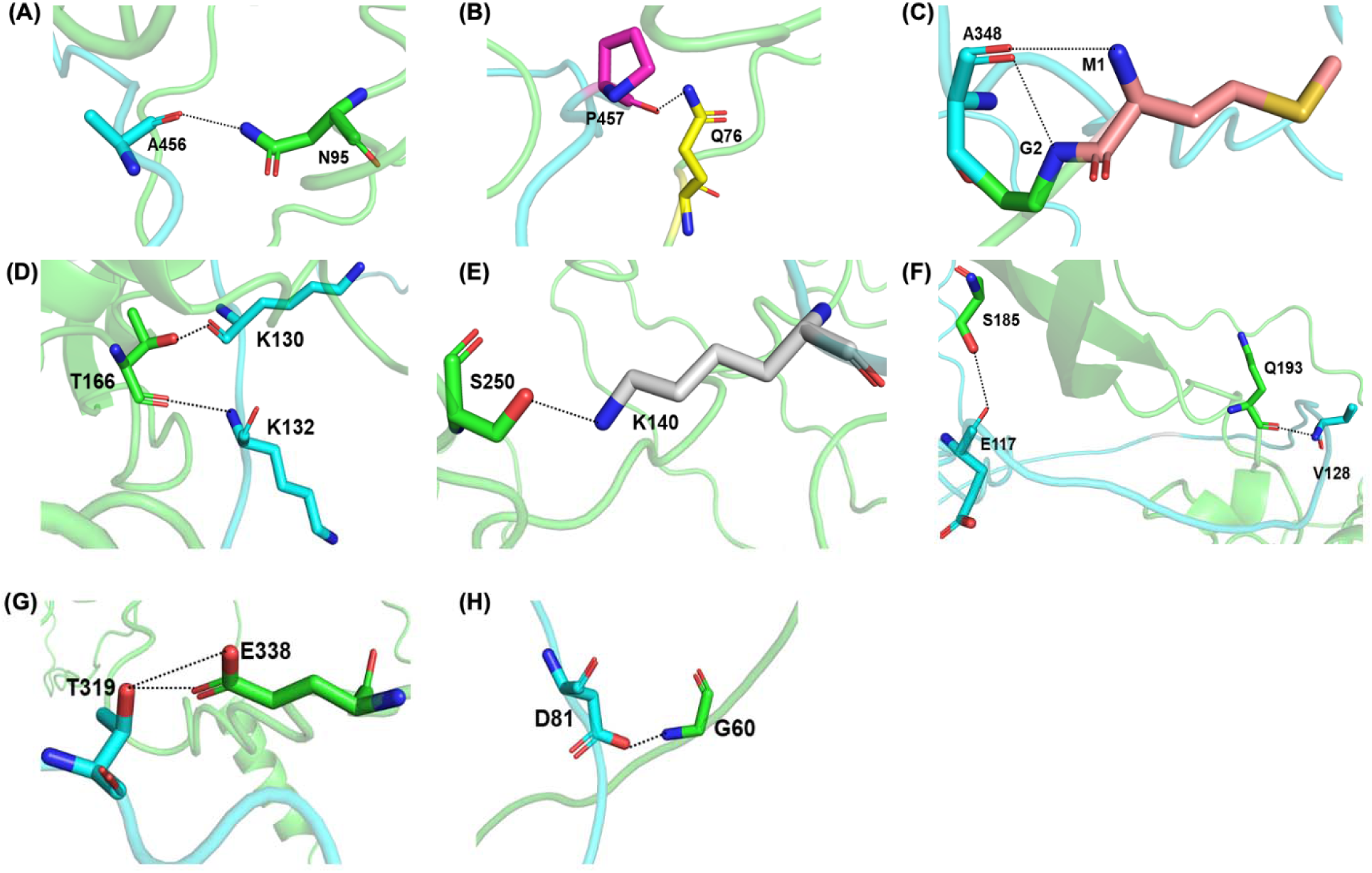
3D visualization of the docked complexes: FG-Nup98-FG-Nup98, FG-Nup98-tau, and tau-tau complexes. Panel (A-C) illustrates the interaction between two FG-Nup98 monomers. Panel (D-F) shows the interaction between an FG-Nup98 monomer and tau. Panel (G-H) depicts the interaction between two tau monomers. In all panels, each monomer is represented in cyan and green, the interacting residues are displayed in stick format, and hydrogen bonds are depicted as black dashed lines.

### Multiscale simulations confirm the superior stability of FG□Nup98 self□association

To comprehend the structural and dynamic behaviours over time, we carried out 100□ns all□atom molecular□dynamics (AAMD) runs and 2□µs coarse□grained simulations (CGMD) for each binary complex. In the CGMD simulations, the elastic network model was utilized due to its computational efficiency and ability to capture essential dynamical features of large biomolecular systems. This approach effectively represented collective motions, functional dynamics, and residue interactions, ensuring a balance between computational cost and biological relevance in simulating protein-protein complexes. Across both simulation scales, the FG□Nup98 homodimer consistently emerged as the most stable species. In the AAMD simulations, the RMSD profiles revealed that the FG-Nup98-FG-Nup98 complex exhibited the highest stability, with an average RMSD of 3.49 nm, while the FG-Nup98-tau complex showed a slightly higher RMSD of 3.92 nm. The tau-tau complex was the least stable, with an average RMSD of 5.50 nm (Figure 6A). CGMD simulations revealed that the FG-Nup98-FG-Nup98 complex demonstrated greater stability, with a lower average RMSD of 0.58 nm. In contrast, the FG-Nup98-tau complex exhibited a higher average RMSD of 3.68 nm, while the tau-tau complex showed the highest RMSD (4.74 nm), indicating reduced stability (Figure 6D). Next, the RMSF analysis showed that the FG-Nup98-FG-Nup98 complex exhibited the average RMSF (1.82 nm in AAMD and 0.48 nm in CGMD), indicating moderate flexibility. The FG-Nup98-tau complex showed lower flexibility with RMSF values of 1.34 nm (AAMD) and 0.43 nm (CGMD). In contrast, the tau-tau complex displayed the highest flexibility, with RMSF values of 1.84 nm (AAMD) and 0.74 nm (CGMD) (Figure 6B, E). Notably, CGMD simulations exhibited lower RMSF values across all systems, reflecting the simplified atomic interactions inherent to this coarse-grained approach. Finally, Rg analysis reinforced the compactness hierarchy. The AAMD simulations indicated that the FG-Nup98-tau and tau-tau complexes retained a more compact structure, whereas the FG-Nup98-FG-Nup98 complex exhibited a lower compactness profile. In AAMD simulations, the average Rg values were 4.15, 4.55, and 4.89 nm for the FG-Nup98-tau, tau-tau, and FG-Nup98-FG-Nup98 models, respectively. Similarly, CGMD simulations yielded corresponding Rg values of 2.85 nm, 3.45 nm, and 2.92 nm (Figure 6C, F). Collectively, the low RMSD, moderate residue□,level fluctuations, and tight Rg of the FG□,Nup98 dimer underscore its structural robustness, while the heterodimer occupies an intermediate regime and higher compactness and tau self□,association is comparatively less compact and dynamic. These multiscale simulation results mirror both the FG-Nup98-tau energetics and our experimental observations.

**Figure 6.**
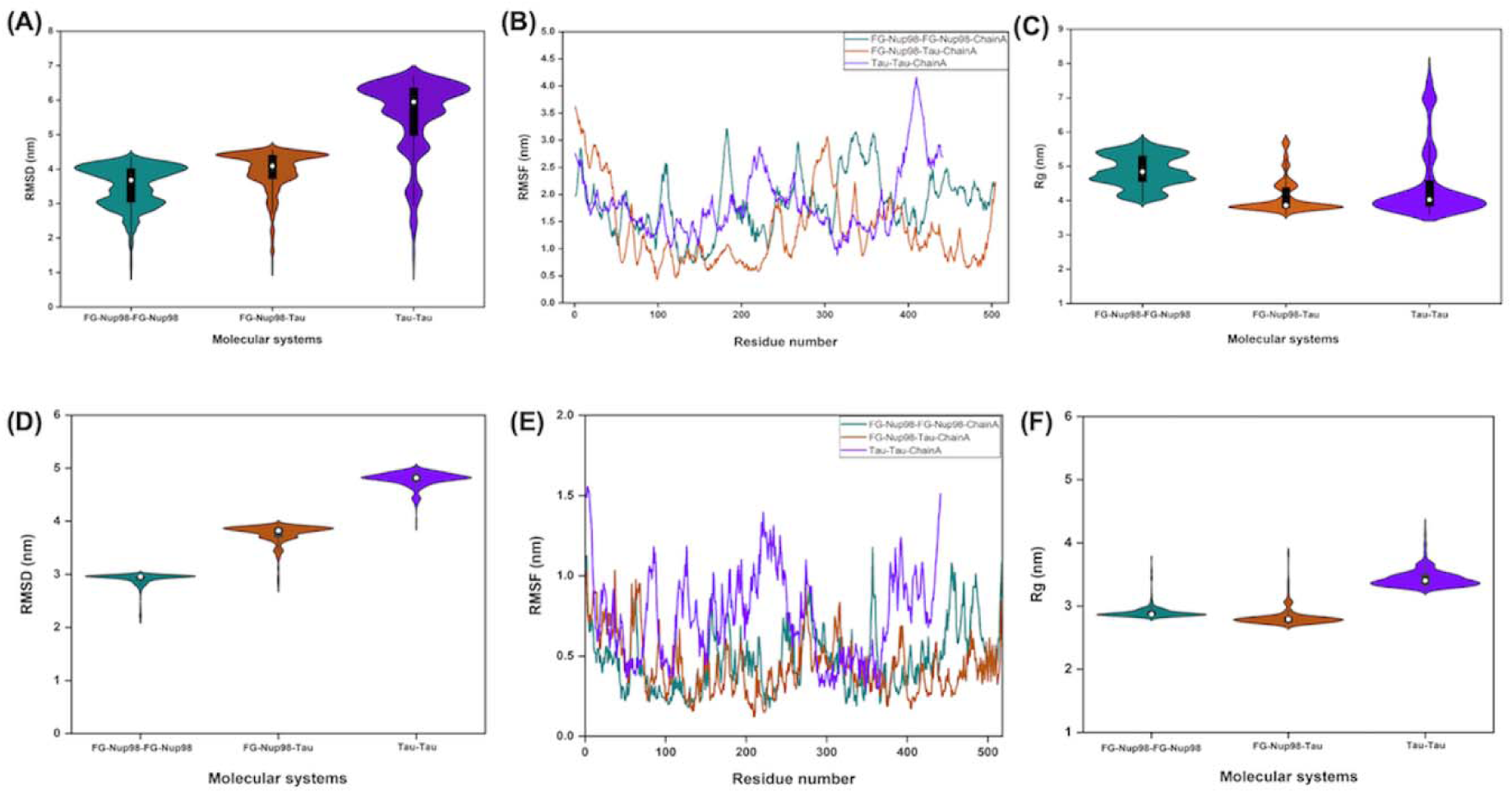
Comparison of the structural stability, flexibility, and compactness FG-Nup98-FG-Nup98, FG-Nup98-tau and tau-tau complexes derived from the AAMD and CGMD simulations. The panels (A-C) show the RMSD, RMSF, and Rg profiles for the FG-Nup98-FG-Nup98, FG-Nup98-tau and tau-tau complexes across 100 ns AAMD simulations. The panels (D-F) depict the RMSD, RMSF, and Rg profiles for the same molecular systems across 2 μs CGMD simulations, each offering insights into stability, fluctuations, and compactness during the long simulations. During the AAMD and CGMD simulations trajectories, the FG-Nup98-FG-Nup98 complex model exhibited the highest stability, minimal inherent fluctuations, and greater compactness compared to the other systems.

### Free□energy landscapes highlight restrained motion of FG□Nup98 homodimers

Principal□component analysis (PCA) was undertaken to describe the dominant, collective motions related to the protein’s secondary structure, sampled by each dimer during the production trajectories. The principal components corresponding to the highest eigenvalues were then used to define the essential subspace, capturing the majority of the dynamic fluctuations. To accomplish this, stable PCA clusters for the FG-Nup98-FG-Nup98, FG-Nup98-tau, and tau-tau complexes were generated using backbone atoms from the equilibrated and stable regions of both AAMD and CGMD trajectories. In AAMD simulations, the covariance matrix trace values were determined as 7924 nm² for FG-Nup98-FG-Nup98, 7318 nm² for FG-Nup98-tau, and 12163 nm² for tau-tau complexes. In contrast, CGMD simulations trace values, measuring 576 nm² for FG-Nup98-FG-Nup98, 905 nm² for FG-Nup98-tau, and 1270 nm² for tau-tau complexes. The first two eigenvectors were the primary contributors to protein dynamics, with Gibbs free energy landscapes (FELs) revealing the most stable states for the FG-Nup98-FG-Nup98 and FG-Nup98-tau complex, as indicated by deeper blue grooves, while the tau-tau complex exhibited greater flexibility and reduced stability, characterized by narrower free energy basins. These FEL profiles were consistent across both AAMD and CGMD simulations (Supplementary Figure S3 A-F), aligning with structural and dynamic variations observed throughout the study. The concurrence of PCA traces, eigenvector projections, and FEL topography across both simulation scales reinforces the earlier dynamical metrics that FG□Nup98□FG□ Nup98 is the most structurally constrained, tau□,FG□, Nup98 is intermediate, and tau□,tau remains the most conformationally promiscuous.

### End□state energy calculations identify tau-tau as the most energetically favoured complex

To quantify how the three dimers are stabilised once solvent effects are taken into account, we extracted equilibrated snapshots from the all□atom trajectories and computed end□state binding free energies with the MM/PBSA protocol, providing insights into the contribution of each energetic term to the overall binding. The equilibrated and stable AAMD simulation trajectories were analyzed to assess binding affinities and elucidate the binding mechanisms for FG-Nup98-FG-Nup98, FG-Nup98-tau, and tau-tau models. The analysis revealed that the homotypic tau-tau complex showed the most favourable binding free energy at −339.64 kcal/mol, surpassing the FG-Nup98-tau and FG-Nup98-FG-Nup98 models, which had binding affinities of −195.45 kcal/mol and −185.73 kcal/mol, respectively (Supplementary Table S2). A detailed decomposition of energetic contributions revealed that ΔVDWAALS, ΔEEL, ΔEGB, ΔGGAS, and ΔSOLV were the primary factors influencing the binding free energies of the complexes. These findings emphasize the greater binding affinity and most favourable energetics of the tau-tau complex compared to the FG-Nup98-tau and FG-Nup98-FG-Nup98 complexes.

### Comprehensive contact analysis underscores the dense molecular interaction network in the tau-tau homodimer

To dissect the molecular underpinnings of the binding free□,energy hierarchy, we analysed equilibrated and stable all□,atom trajectories, cataloguing every hydrogen bond, hydrophobic contact, polar and ionic interaction, van der□,Waals contact, aromatic stacking, and carbonyl contact formed across each interface. The tau-tau complex emerged as the most interaction-rich assembly, with a total of 2205 interactions. This was followed by 1927 interactions for the FG-Nup98-tau heterodimer and 1561 interactions for the FG□,Nup98-FG-Nup98 self-association. The sheer abundance and diversity of contacts in the homodimer, including spanning hydrogen bonding, electrostatic, and non□,polar interactions, provide a structural rationale for its superior MM/PBSA binding free energy and reinforce the experimental picture of tau-tau acting as a multivalent partner that avidly recruits tau (Supplementary Table S3).

## Discussion

In this work, we have elucidated the molecular basis of co-condensation between the FG-repeat domain of the nucleoporin Nup98 and the microtubule-associated protein tau, and determined how stoichiometry, environment, and intrinsic interactions dictate condensate architecture and dynamics. These findings provide mechanistic insight into a putative early stage of tauopathy, in which FG-Nup98-tau assemblies could impair NCT and seed aberrant tau aggregation. We now discuss three major themes emerging from our data: (i) the sequence-driven LLPS behavior of FG-Nup98 and tau and their simulation-derived interaction hierarchy, (ii) the stoichiometry- and environment-dependent architecture and material properties of heterotypic condensates, and (iii) the potential implications for tau-mediated NCT disruption and therapeutic strategies.

Our bioinformatic analyses established that both full-length tau and the FG domain of Nup98 are highly intrinsically disordered, present high predicted pLLPS scores (0.9951 for tau and 1.000 for FG-Nup98), and exhibit charged, low-complexity features consistent with multivalent weak interactions. This is in line with prior studies showing tau to be intrinsically disordered, with flexible N-terminal and repeat domains that mediate aggregation and interactions ^23, 24, 25^, and FG-Nups to be classic IDPs that form the permeability barrier through weak hydrophobic/FG-FG contacts ^9, 11, 26, 27^. The fact that both partners have high intrinsic LLPS propensities sets the stage for heterotypic condensation: that is, FG-Nup98 and tau are biophysically poised to co-assemble, rather than simply aggregate. The docking and simulation studies provide further atomic□,level substantiation of multivalency: FG-Nup98-FG-Nup98 interactions yield the strongest docking scores, followed by FG-Nup98-tau, and then tau-tau. Although tau-tau contacts are frequent in the simulations, FG-Nup98 provides the stronger and more structured network of interactions, giving it a scaffold-like role, which recruits tau into condensates. This hierarchical affinity landscape echoes recent work on IDP condensates in neurodegeneration, where scaffold and client relationships determine organization and composition of droplets ^18^. Moreover, the ability of FG-Nup98 to self□,associate (highest affinity) suggests that in cells, FG□repeat domains may form an inner core that then sequesters tau as a client, rather than a simple co□mixing.

The simulation data reinforce this view: the all□,atom and coarse□,grained MD results show that FG-Nup98 homodimers have the lowest RMSD, lowest RMSF, tightest Rg, and deepest free□,energy basins. By contrast, tau-tau assemblies are more dynamic and less compact, but enjoy more favorable MM/PBSA binding free energies. Interestingly, though perhaps counterintuitive, the stronger binding energy of tau-tau might reflect many more weak interactions, and we propose that in the context of heterotypic droplet assembly, the relative kinetics, concentrations, and multivalent scaffolding likely shift the equilibrium toward an FG-Nup98-dominated architecture. In sum, the sequence, docking, and simulation data provide a consistent molecular rationale for why and how FG-Nup98 can recruit tau into heterotypic condensates.

A central finding of our study is that the stoichiometry of FG-Nup98-tau strongly governs the architecture of the condensates. When tau is in large excess (≥ 10:1), droplets are homogenously mixed and both proteins colocalize in the same dense phase. At more balanced stoichiometries (5:1 to 2:1), a mixed population emerges: colocalized droplets co-exist with “nested” droplets in which FG-Nup98-rich subdomains sit inside larger tau□rich droplets. At 1:1 or inverted stoichiometry, isolated FG-Nup98 droplets (with or without tau) or nested architectures predominate, and at high FG-Nup98 concentration, aggregates form. This tunable organization is reminiscent of prior observations of scaffold–client systems and sub□compartmentalized condensates (droplets within droplets) in other IDP systems ^18, 28^. The fact that stoichiometry controls demixing vs mixing suggests that the relative valency and concentration of FG-Nup98 vs tau determine whether the system forms a single dense phase or phase□separated sub-phases. From a physical perspective, when tau dominates, its weaker tau-tau and FG-Nup98-tau contacts suffice to maintain a single mixed dense phase; when FG-Nup98 concentration increases, higher FG-Nup98-FG-Nup98 affinity drives internal segregation, forming FG-rich cores inside tau shells.

The environmental modulation experiments further demonstrate the sensitivity of these condensates to macromolecular crowding (PEG) and ionic strength. Under 10% PEG and low salt (0 mM NaCl), the droplets are uniformly mixed and highly liquid□,like; at physiological salt (∼150 mM), demixing appears, and at high salt (500 mM to 1 M), the droplets collapse into amorphous clusters or FG-Nup98-only aggregates. This behaviour aligns with the known salt□sensitivity of LLPS of tau (inhibited by high salt) and the promoting effect of salt on FG□Nup98 LLPS (which arises from hydrophobic FG interactions) ^19, 20, 21, 22^. Taken together, these findings imply that subtle changes in local concentration, ionic strength, or crowding (all of which could vary in a stressed neuron) may tip FG-Nup98-tau assemblies between liquid□like, homogeneous droplets on one hand and nested, segregated (or ultimately aggregated) structures on the other. Importantly, the nested architecture (FG-Nup98 cores embedded in tau shells) is reminiscent of “droplet within droplet” morphologies seen in other multicomponent condensates ^18, 28^, and suggests a route by which tau may become locally enriched in an FG-Nup scaffolded droplet, which enhances its local concentration and perhaps its propensity for misfolding or fibrillization.

The material properties of the condensates were probed via FRAP. In mixed droplets, tau recovers ∼34% and FG-Nup98 ∼48% of the pre□,bleach signal, which indicates moderate mobility of both components. However, in nested droplets, tau recovery drops to ∼21% and the FG□,Nup98 core is essentially immobile (unable to bleach). This suggests that in nested assemblies, the FG core acts as a quasi□,solid scaffold that severely restricts tau diffusion. This finding has two major implications: first, the nested architecture correlates with decreased internal dynamics, and second, such decreased mobility may favour recruitment of additional tau molecules (or other binding partners) and slower turnover, which potentially increases the local residence time and promotes pathological conversion. In the cellular context, this means that when FG-Nup98/tau stoichiometry or environment favours nested droplet formation, tau may become kinetically trapped and accumulate in restricted sub□,regions. This means a scenario that could promote nucleation of tau fibrils or block dynamic NCT.

Our findings bear important implications for understanding how tau may disrupt the NPC and NCT in tauopathies such as AD. Accumulating evidence shows that mislocalised tau (especially phosphorylated tau) interacts with NPC components, including Nup98, alters nuclear envelope morphology, and impairs NCT. For instance, Eftekharzadeh et al. demonstrated that tau interacts with Nups and disrupts the process of NPC ^4^. A recent review by Diez & Wegmann summarises the mechanisms of tau-nucleus interactions and suggests that tau-Nup binding may precede fibril formation in disease ^29^. Our work offers a biophysical mechanism for this: the heterotypic phase separation of FG□,Nup98 with tau yields condensates that could physically sequester tau in proximity to NPCs or Nups, reduce mobility within the droplet, and thereby favour dysfunctional accumulation. In the nested architecture, the FG core acts like a scaffold that traps tau and limits its diffusional exit, which may equate to impaired release of tau from NPC complexes or inhibited recycling of transport factors. Moreover, the stoichiometry□,dependent architecture suggests that when FG-Nup98 expression is altered (for example, mislocalised or over-expressed in disease), or when tau concentration rises (due to aggregation-prone isoforms or stress□,induced misfolding), the balance may shift to nested or aggregated droplet forms that are more rigid and less dynamic. That could correspond to early events in disease progression, where Nup-tau condensates form, trap transport factors or NPC components, and seed further tau aggregation. The high local concentration of tau in such condensates could accelerate its conversion to oligomers or fibrils. Indeed, previous *in vitro* studies show that FG-Nup98 accelerates tau fibrillization ^8^. From the perspective of transport□,mechanism, FG-Nup condensates normally act as the selective barrier for nuclear import/export, with disordered FG domains forming a dynamic mesh (selective phase model) ^30^. When tau enters this system and forms a heterotypic phase, several paths of dysfunction emerge: (a) Tau may displace or perturb FG-Nup-FG-Nup interactions, which alters barrier permeability; (b)FG-Nup-tau condensates may reduce the mobility of transport receptors or cargo; (c) nested/aggregated FG-Nup-tau condensates may physically clog or sequester NPCs, which reduces transport flux; or (d) tau may promote transition from liquid to gel/solid states of FG-Nups that further stiffens the barrier. Our observations that high salt (which mimics cellular stress) shifts the interaction toward aggregated FG-Nup clusters suggest that stressed neuronal conditions may move the system from functional to dysfunctional states. Given that nuclear transport deficits are now being widely appreciated in neurodegenerative diseases, including ALS, FTD, and AD, and that FG-Nup malfunction is implicated in these, our nested□,droplet model offers a plausible mechanistic link between the interaction of FG-Nup98 and tau and the NCT disruption, which may inform future therapeutic strategies ^31^.

Our study integrates *in vitro* droplet assays, advanced microscopy (including FRAP), and multiscale molecular simulations. However, some caveats and limitations merit discussion. First, the *in vitro* droplet system uses recombinant proteins in simplistic buffer/crowder setups; cellular environments include chaperones, other Nups, nuclear transport receptors, and protein PTMs that may modulate behaviour. For example, the phosphorylation of tau reportedly promotes accumulation with Nup98, while oligomerisation of tau reduces Nup98 binding ^8^. Future work should test the influence of PTMs, additional Nups, and transport receptors. Second, while docking and simulations provide mechanistic insight, they are performed on monomeric models rather than within highly multivalent droplet environments; the actual network of interactions in condensates is likely richer and more dynamic. Third, the droplet□within□droplet (“nested”) architecture is an *in vitro* observation; direct evidence for such nested condensates in neurons or at NPCs remains to be obtained. Correlative microscopy of NPCs and tau in brain tissue would strengthen the physiological relevance. Nonetheless, the tunable architecture and data for material properties provide a quantitative framework that bridges molecular interactions and mesoscale condensate behaviour.

## Conclusions and perspectives

Conceptually, our findings contribute to two growing themes in cell biology: the role of multicomponent phase separation in cellular organization, and the emerging link between LLPS and neurodegeneration. The idea that heterotypic condensates (rather than homotypic ones) can yield nested sub-compartments is increasingly recognized. Here, we show how stoichiometry and interaction hierarchy (scaffold/ client) drive internal architecture. Also, the link between tau and an NPC component via LLPS suggests that mislocalised IDPs may exploit existing condensate scaffolds (such as FG-Nups) to seed pathological assemblies. Thus, phase separation may not merely be a by-product of aggregation, but a mechanistic gateway from physiological dynamic condensate to pathological rigid organelle.

From a translational perspective, our mechanistic framework suggests several possible entry points for intervention in tauopathy. If the early step is tau recruitment into FG-Nup condensates, then strategies aimed at stabilising the functional, liquid□like state of FG-Nup condensates, or reducing tau recruitment, could preserve NCT and prevent subsequent aggregation. For example, small molecules or peptides that disrupt FG□Nup-tau interactions, or modulate FG network dynamics (e.g., via importins or small modulators of FG–FG binding) may be protective. Indeed, importin binding has been shown to suppress LLPS of other IDPs ^32^. Moreover, modulating the intracellular environment (via osmolytes, ionic channels, or stress pathways) might shift the droplet behaviour back toward the functional regime. At the level of diagnosis, nested or rigid FG-Nup-tau condensates could represent early biomarkers of NCT dysfunction, visible before tau fibrillation. From a research perspective, key next steps include (i) validating nested condensate architecture in neurons or brain tissue, (ii) probing how tau PTMs (phosphorylation, acetylation, truncation) alter the stoichiometry–architecture map, (iii) examining how other NPC components (e.g., Nup62, Nup153) participate in tau recruitment, (iv) using time-resolved live-cell imaging of FG-Nup98-tau condensate formation in stressed neurons, and (v) screening for modulators of FG-Nup98-tau condensation.

In summary, we have presented a mechanistic dissection of heterotypic phase separation between the FG-repeat domain of Nup98 and tau, and demonstrated how stoichiometry, crowding, and ionic strength dictate condensate architecture and material behaviour, and how the relative binding hierarchies and dynamics drive recruitment of tau into FG-Nup scaffolds. These insights not only deepen our molecular understanding of how tau may impact the nuclear pore and NCT but also highlight new mechanistic pathways by which phase separation may shift from physiological to pathological states in neurodegeneration. Our study thus bridges the mesoscale world of biomolecular condensates with the molecular pathology of tauopathy, and points toward novel strategies to intercept the early events of nuclear transport dysfunction and tau aggregation.

## Methods

### Bioinformatic analysis

The protein sequences of full-length human tau (UniProt ID: P10636-8) and the 3Cys-mutant of the FG-domain of human Nup98 (UniProt ID: P52948) were used for bioinformatics analysis. Disorder propensity was predicted using the RIDAO meta-server, which integrates six prediction tools ^15^. Net Charge Per Residue (NCPR) was calculated using the CIDER platform (https://pappulab.wustl.edu/CIDER/) ^16^. Phase separation propensities were estimated using the FuzDrop server (https://fuzdrop.bio.unipd.it/), which reports pLLPS scores based on sequence features and known LLPS drivers ^17^.

### Overexpression and purification of FG-Nup98

To assess the expression of FG-Nup98, the recombinant *FG-Nup98*-pPEP-TEV construct, which was a kind gift from Dr. Roderick Lim (University of Basel, Switzerland), was transformed into *E. coli* BL21 cells and plated on LB agar containing ampicillin (100 µg/mL). A single colony was inoculated into 5 mL of LB broth supplemented with 100 µg/mL ampicillin and grown overnight at 37°C with continuous shaking. This starter culture was used to inoculate 500 mL of LB medium with ampicillin (100 µg/mL) in a 1:100 ratio and incubated with shaking at 37°C until the OD□□□ reached approximately 0.6-0.8. The culture was cooled, induced with 0.5 mM isopropyl β-D-1-thiogalactopyranoside (IPTG), and incubated further at 18°C with shaking at 130 rpm for 22 hours. The cells were harvested by centrifugation at 4°C and stored at −80°C until further use.

The protein was purified using a protocol partially adapted from Kowalczyk *et al.,* 2011 ^33^, with modifications. For lysis, the cells were resuspended in ∼15 mL of equilibration buffer, henceforth referred to as buffer A (10 mM Tris-Cl, pH 8, 100 mM NaCl, 6 M urea). A 1X protease inhibitor cocktail was added, and the cells were lysed on ice using sonication (50 Hz amplitude; 30-second pulse with 30-second interval; 30 cycles). The crude lysate was clarified by centrifugation at 12,000 rpm for 30 minutes at 4°C, and the supernatant was filtered through a 0.45 µm polyvinylidene difluoride (PVDF) membrane. Purification was performed using Ni²□-NTA affinity chromatography. The column matrix was washed extensively with 10 bed volumes of buffer A, and the supernatant was applied to the matrix. Washing was carried out using increasing concentrations of imidazole, and the recombinant protein was eluted with elution buffer containing 10 mM Tris-Cl (pH 8), 100 mM NaCl, 6 M urea, and 300 mM imidazole, collected in two fractions of 2 mL each. The purified protein was analyzed on a 12% SDS-PAGE gel and dialyzed at 4°C in dialysis buffer (20 mM phosphate buffer, pH 7.4, 100 mM NaCl, 4mM DTT) to remove imidazole and excess urea. The protein was filtered through a 0.22 µm PVDF membrane, and the concentration was determined spectrophotometrically using the extinction coefficient of FG-Nup98, 5960 M□¹ cm□¹. The protein was stored at −20°C until further use.

### Overexpression and purification of Tau

To assess the expression of tau, the recombinant *tau*-pET29b construct was transformed into *E. coli* BL21 cells and plated on LB agar containing kanamycin (50 µg/mL). A single colony was inoculated into 5 mL of LB broth supplemented with 50 µg/mL kanamycin and grown overnight at 37°C with continuous shaking. This starter culture was used to inoculate 500 mL of LB medium with kanamycin (50 µg/mL) in a 1:100 ratio and incubated with shaking at 37°C until the OD□□□ reached approximately 0.6-0.8. The culture was induced with 0.5 mM IPTG and incubated further at 37°C with shaking at 180 rpm for 4 hours. The cells were harvested by centrifugation at 4°C and stored at −80°C until further use.

The protein was purified using a protocol adapted from Poudyal *et al*., 2022 ^34^. The cells were resuspended in ∼15 mL of equilibration buffer, henceforth referred to as buffer B, containing 20 mM phosphate, pH 7.4, and 50 mM NaCl, for cell lysis. A 1X protease inhibitor cocktail and 4 mM DTT were added, and the cells were lysed on ice using sonication (40 Hz amplitude; 3-second pulse with 3-second interval; 15 cycles). The crude lysate was then subjected to boiling at 95°C for 20 minutes, after which the solution was clarified by centrifugation at 12,000 rpm for 30 minutes at 4°C. 10% streptomycin and glacial acetic acid were added to the supernatant, and kept at 4°C with slow rocking for 30 minutes. The solution was centrifuged again at 12,000 rpm for 30 minutes at 4°C. To the supernatant, an equal volume of saturated ammonium sulphate was added and kept at 4°C, with slow rocking overnight. The solution was then centrifuged again at 12,000 rpm for 30 minutes at 4°C, and the pellet was retrieved. The pellet was dissolved in 100 mM ammonium acetate and reprecipitated with 100% ethanol, and the solution was centrifuged at 12,000 rpm for 30 minutes at 4°C. This step was repeated three times. The final pellet was dissolved in ∼2 mL of 20 mM phosphate, pH 7.4, 50 mM NaCl, and 4 mM DTT. The protein was filtered through a 0.22 µm PVDF membrane, and the concentration was determined spectrophotometrically using the extinction coefficient of tau, 7575 M□¹ cm□¹. It was also analyzed on a 12% SDS-PAGE gel. The protein was stored at −20°C until further use.

### Fluorescent labeling of proteins

Both proteins were labeled using thiol-active fluorescent AlexaFluor dyes: tau labeled with AlexaFluor 488 C5 maleimide, and FG-Nup98 using AlexaFluor 594 C5 maleimide. The labeling process was conducted using an established protocol ^35^. For fluorescent labeling of tau, the AlexaFluor 488 C5 maleimide dye was dissolved in dimethyl sulfoxide (DMSO) and mixed with the protein at a 1:10 ratio, with continuous stirring. The reaction mixture was incubated at 20°C for 20 minutes. Then, the protein sample was incubated at 4 °C for 24 hours, with slow mixing. Excess free dye was removed by extensive dialysis in sodium phosphate buffer (pH=7.4). For FG-Nup98, the same process was carried out with the AlexaFluor 594 C5 maleimide dye.

### *In vitro* protein phase separation experiments using spectrophotometry and confocal microscopy

The initial phase separation screening was performed using turbidity assay measurements following the protocol by Ferrolino *et al*., 2018 ^36^. A spectrophotometer (Thermo Scientific Genesys) was utilized to assess sample turbidity at an excitation wavelength of 340 nm. To confirm the phase separation of FG-Nup98 and Tau, droplet imaging was conducted using confocal microscopy. For these experiments, 10% labelled protein was mixed with unlabelled protein. Following 1 hour of incubation, the protein samples were transferred onto a grease-free, thoroughly cleaned 22 mm cover glass (Blue-Star, India). Confocal microscopy was performed on an inverted Leica TCS SP8 confocal microscope (Germany) equipped with a 488 nm laser to excite Alexa 488 maleimide-labelled tau and a 561 nm laser to excite Alexa 594 maleimide-labeled FG-Nup98. Imaging was carried out with a 63× (1.4 NA) oil immersion objective lens. The captured images had a resolution of 1024 × 1024 pixels, with a line averaging of 3 and a frame accumulation of 1. All confocal images were analysed and processed using ImageJ software (National Institutes of Health, USA).

### Fluorescence recovery after photobleaching (FRAP)

FRAP was used to assess the material properties of droplets *in vitro*. The imaging was performed using an inverted TCS SP8 confocal microscope (Leica). Protein samples were bleached using a 488 nm laser set to 80% power. The experimental parameters included a pre-bleach duration of 3 seconds, a bleach duration of 20 milliseconds, and a recovery period of 120 seconds. For these measurements, a region of interest (ROI) with a diameter of 1.5 μm was selected from the images.

### System preparation for molecular docking and simulation experiments

Since the monomeric structures of FG-Nup98 and tau were unavailable, we predicted them using AlphaFold v2.0. The tau (UniProt ID: P10636-8) sequence was retrieved from the UniProt, and the sequence for the 3Cys-mutant of the FG-domain of human Nup98 (UniProt ID: P52948) was provided by Dr. Roderick Lim (University of Basel, Switzerland), which was used as input for structure prediction. Specifically, monomeric FG-Nup98 and tau structures were generated using AlphaFold v2.0 ^37, 38^, employing the run_docker.py Python script. This process generated five models, from which the top-ranked structure was selected for further analysis based on confidence scores and predicted local distance difference test (pLDDT) scores. The selected monomeric structures were subsequently used for protein-protein interaction studies.

### Protein-protein docking of FG-Nup98-FG-Nup98, FG-Nup98-tau, and tau-tau monomers

To investigate the key biological functions, intermolecular interactions, and binding affinities of FG-Nup98 and tau monomers, particularly their interactions in FG-Nup98-tau, FG-Nup98-FG-Nup98, and tau-tau complexes, we performed a series of protein-protein docking studies using the LZerD algorithm ^39^. The docking studies were performed with the monomeric structures of FG-Nup98 and tau as initial models. Utilizing the LZerD algorithm, we analyzed the binding orientations and conformations of FG-Nup98-tau, FG-Nup98-FG-Nup98, and tau-tau complexes, comparing the predicted interactions with experimentally reported data. Each docking experiment produced multiple binding conformations, from which the optimal models were chosen based on the GOAP score, DFIRE score, ITScore score, Ranksum Score, and number of intermolecular interactions formed. The Ranksum score, representing the sum of the ranks from the three scores, is primarily used for ranking the quality of the model, with lower scores suggesting better binding affinities. To gain deeper insights, PyMOL (http://www.pymol.org/) was used to visualize the docked complexes, their binding orientations, conformations, and polar contacts, enabling a detailed analysis of the binding interfaces and intermolecular interactions.

### Insights into the structural dynamics of FG-Nup98-FG-Nup98, FG-Nup98-tau, and tau-tau complexes through AAMD and CGMD simulations

To investigate various physicochemical properties such as the structural dynamics, stability, and compactness of the FG-Nup98-FG-Nup98, FG-Nup98-tau, and tau-tau complexes, we employed both all-atom molecular dynamics (AAMD) and coarse-grained molecular dynamics (CGMD) simulations. While docking studies provide a static representation of binding interactions, they fail to capture the dynamic behavior of such protein-protein complexes. To overcome this limitation, we performed AAMD and long-timescale CGMD simulations to assess various physicochemical properties and binding mechanisms of FG-Nup98-FG-Nup98, FG-Nup98-tau, and tau-tau interactions over time.

### All-atom molecular dynamics simulations

Using the CHARMM36 force field ^40^ and the GROMACS v.2023 package ^41, 42^, we simulated three systems: FG-Nup98-FG-Nup98, FG-Nup98-tau, and tau-tau complexes. Following this, each system was solvated using a TIP3P water model in a cubic box, ensuring a minimum distance of 1.0 nm from the box edge. To neutralize the system charges, 16 Cl− ions were added to the FG-Nup98-FG-Nup98 complex, 10 Cl− ions to FG-Nup98-tau, and 4 Cl− ions to the tau-tau complex. Energy minimization was carried out using the steepest descent algorithm for 50,000 steps while preserving periodic boundary conditions. Next, the systems underwent a heating phase in the canonical (NVT) ensemble, followed by equilibration in the isothermal-isobaric (NPT) ensemble. For each system, production runs were conducted in the NPT ensemble for 100 ns. Important simulation parameters included the use of the Parrinello-Rahman barostat ^43^ to maintain constant pressure, the Berendsen thermostat ^44^ for temperature coupling, and a 2 fs integration time step to ensure efficient time evolution. A Coulomb interaction cutoff of 1.2 nm and a Fourier grid spacing of 0.16 nm were applied. Long-range electrostatics were handled using the Particle Mesh Ewald (PME) method ^45^, while the LINCS algorithm ^46^ was employed to constrain bond lengths. This simulation framework offered valuable insights into the structural dynamics and stability of the FG-Nup98-FG-Nup98, FG-Nup98-tau, and tau-tau complexes.

### Coarse-grained molecular dynamics simulations

After AAMD simulations, we conducted coarse-grained simulations ^47^ for the FG-Nup98-FG-Nup98, FG-Nup98-tau, and tau-tau complexes. Coarse-grained simulations provide improved computational efficiency, enabling the exploration of longer timeframes and larger length scales. Using the MARTINIZE2 tool, we converted a refined atomistic protein structure into a coarse-grained (CG) model and generated the corresponding topology file with the Martini2.0 force field ^48^. The secondary structure was assigned using the -dssp flag ^49^, while the elastic network model applied harmonic constraints between atom pairs to simulate the connectivity and flexibility of the FG-Nup98-FG-Nup98, FG-Nup98-tau, and tau-tau complexes. Each system was prepared using the insane.py script, which included solvation and ion addition for neutralization. Energy minimization was then carried out using the steepest descent algorithm for 50,000 steps to stabilize the system. Following system preparation, a 50 ns equilibration run was performed. After confirming system convergence, 2 μs production simulations were conducted for each system, totalling 6 μs. The simulations were carried out under NPT ensemble conditions with periodic boundary conditions, employing modified Berendsen temperature coupling and maintaining a constant pressure of 1 atm. This method ensured stable and realistic dynamic behavior of the FG-Nup98-FG-Nup98, FG-Nup98-tau, and tau-tau complexes throughout the simulations, maintaining structural integrity and physiologically relevant interactions.

### Analysis of various physicochemical properties and molecular interactions from AAMD and CGMD simulation trajectories

The physicochemical properties of the FG-Nup98-FG-Nup98, FG-Nup98-tau, and tau-tau models were evaluated using AAMD and CGMD simulation trajectories. Structural stability, flexibility, and compactness were assessed through root-mean-square deviation (RMSD), root-mean-square fluctuations (RMSF), and the radius of gyration (Rg), respectively. These analyses were performed using GROMACS with the gmx_mpi rms, gmx_mpi rmsf, and gmx_mpi gyrate modules, respectively, to track structural deviations over time.

Additionally, molecular interactions were analyzed from stable AAMD simulations using the gmx_mpi hbond module and the Arpeggio server ^50^ to assess binding characteristics. Next, principal component analysis (PCA) was conducted to examine major conformational changes and collective motions ^51^. A variance-covariance matrix was constructed from atomic coordinates, and eigenvectors and eigenvalues were extracted using the gmx_mpi covar and gmx_mpi anaeig modules in GROMACS. The dominant modes of motion were identified by projecting simulation trajectories onto the first principal components, providing deeper insights into system dynamics. Additionally, the free-energy landscape (FEL) was constructed to analyze conformational stability and energy states throughout the simulations. The gmx_mpi sham module was utilized to generate 3D contour plots, depicting conformational energy minima and offering insights into the stability and dynamic equilibrium of the FG-Nup98-FG-Nup98, FG-Nup98-tau, and tau-tau complexes throughout the simulations.

### MM/PBSA-based binding free-energy estimation from AAMD trajectories

The end-state binding free energies for the FG-Nup98-FG-Nup98, FG-Nup98-tau, and tau-tau complexes were computed from AAMD simulation trajectories using the MM/PBSA-based approach. These calculations were performed with the gmx_mmpbsa tool, which incorporates the MMPBSA.py script from AMBER for binding energy estimation. The setup was implemented in a Miniconda environment using Python 3, ensuring all necessary dependencies for gmx_mmpbsa were properly integrated ^52, 53^. The analysis was conducted using various input files, including the MMPBSA Python script, trajectory file (.xtc), topology file (.top), and index files defining FG-Nup98-FG-Nup98, FG-Nup98-tau, and tau-tau as separate groups. Additionally, the protein structure file (.pdb), GROMACS topology file (.gro), and portable topology file (.tpr) were utilized for the calculations. The binding free energy calculations were conducted using stable and equilibrated AAMD trajectory data, ensuring analysis was based on a period of stable simulation behavior.

## Data availability

All data supporting the findings of this study are available within the manuscript and the supplementary information.

## Acknowledgements

T.T. would like to acknowledge the support of a project grant from the Indian Council of Medical Research (ICMR), Government of India, India (52/06/2020/BIO/BMS). N.N. would like to thank DST-INSPIRE for providing fellowship. The authors also acknowledge the infrastructure facilities of IIT (BHU) Varanasi and DST-funded I-DAPT Hub Foundation, IIT (BHU) [DST/NMICPS/ TIH11/IIT(BHU)2020/02]. Further, the computing resources of PARAM Shivay Facility under the National Supercomputing Mission, Government of India, at the IIT (BHU), Varanasi, are gratefully acknowledged.

## Contributions

Conceptualization, TT, KC, NN; Methodology, NN, SR, AJY; Validation, NN, SR, AJY, AKP; Formal analysis, NN, SR, AJY; Investigation, NN, SR, AJY; Writing—original draft preparation, NN, AJY; Writing—review and editing, TT, KC, AKP; Supervision, TT, KC; Funding acquisition, TT.

## Competing interests

The authors declare no competing interests.

## Notes

### Competing Interest Statement

The authors have declared no competing interest.

