## Supplementary files for "Heterotypic FG-Nup98-tau condensates form nested assemblies through stoichiometry-dependent phase transitions"

**Supplementary Table S1**. Summary of molecular systems used in docking experiments with LZerD algorithm, including their respective docking scores (kcal/mol).

| **Molecular systems** | **GOAP Score** | **DFIRE Score** | **ITScore score** | **Ranksum Score** |
| --- | --- | --- | --- | --- |
| FG-Nup98-FGNup98 complex | -25270.73 | -25724.44 | -12118.18 | 17 |
| FG-Nup98-Tau complex | -18989.09 | -19681.47 | -9188.73 | 33 |
| Tau-Tau complex | -12711.30 | -13690.87 | -6086.16 | 19 |

**Supplementary Table S2**. The table presents the end-state binding free energy values (in kcal/mol) from stable AAMD simulation trajectories of tau-tau, FG-Nup98-tau, and FG-Nup98-FG-Nup98 complexes calculated using the MM/PBSA-based method.

| **Molecular systems** | **Energy component** | **Average** | **SD**  ${\boldsymbol{(}\mathbf{Prop}\mathbf{.)}}^{\boldsymbol{a}}$ | ${\boldsymbol{(}\mathbf{SD}\mathbf{)}}^{\boldsymbol{b}}$ | **SEM**  ${\boldsymbol{(}\mathbf{Prop}\mathbf{.)}}^{\boldsymbol{c}}$ | ${\boldsymbol{(}\mathbf{SEM}\mathbf{)}}^{\boldsymbol{d}}$ |
| --- | --- | --- | --- | --- | --- | --- |
| **Tau-Tau** | ΔBOND | 0 | 17.1 | 0 | 1.21 | 0 |
|  | ΔANGLE | 0 | 27 | 0 | 1.9 | 0 |
|  | ΔDIHED | 0 | 14.42 | 0 | 1.02 | 0 |
|  | ΔUB | 0 | 5.23 | 0 | 0.37 | 0 |
|  | ΔIMP | 0 | 6.83 | 0 | 0.48 | 0 |
|  | ΔCMAP | 0 | 13.81 | 0 | 0.97 | 0 |
|  | ΔVDWAALS | -378.04 | 13.25 | 44.46 | 0.93 | 3.14 |
|  | ΔEEL | -4351.18 | 13.32 | 339 | 0.94 | 23.91 |
|  | Δ1-4 VDW | 0 | 7.93 | 0 | 0.56 | 0 |
|  | Δ1-4 EEL | 0 | 44.88 | 0 | 3.17 | 0 |
|  | ΔEPB | 4447.89 | 42.98 | 340.33 | 3.03 | 24 |
|  | ΔENPOLAR | -58.31 | 3.56 | 6.08 | 0.25 | 0.43 |
|  | ΔGGAS | -4729.22 | 24.85 | 367.38 | 1.75 | 25.91 |
|  | ΔGSOLV | 4389.58 | 43.13 | 335.68 | 3.04 | 23.68 |
|  | **ΔTOTAL** | **-339.64** | **49.78** | **39.1** | **3.51** | **2.76** |
| **FG-Nup98-Tau** | ΔBOND | 0 | 18.94 | 0 | 1.34 | 0 |
|  | ΔANGLE | 0 | 25.44 | 0 | 1.79 | 0 |
|  | ΔDIHED | 0 | 13.81 | 0 | 0.97 | 0 |
|  | ΔUB | 0 | 4.42 | 0 | 0.31 | 0 |
|  | ΔIMP | 0 | 6.24 | 0 | 0.44 | 0 |
|  | ΔCMAP | 0 | 10.24 | 0 | 0.72 | 0 |
|  | ΔVDWAALS | -341.85 | 17.58 | 22.04 | 1.24 | 1.55 |
|  | ΔEEL | -294.02 | 70.71 | 80.82 | 4.99 | 5.7 |
|  | Δ1-4 VDW | 0 | 8.81 | 0 | 0.62 | 0 |
|  | Δ1-4 EEL | 0 | 40.66 | 0 | 2.87 | 0 |
|  | ΔEPB | 485.73 | 38.75 | 86.09 | 2.73 | 6.07 |
|  | ΔENPOLAR | -45.3 | 1.67 | 2.55 | 0.12 | 0.18 |
|  | ΔGGAS | -635.87 | 73.98 | 93.08 | 5.22 | 6.57 |
|  | ΔGSOLV | 440.43 | 38.79 | 84.42 | 2.74 | 5.95 |
|  | **ΔTOTAL** | **-195.45** | **83.53** | **18.61** | **5.89** | **1.31** |
| **FG-Nup98-FG-Nup98** | ΔBOND | 0 | 19.04 | 0 | 1.34 | 0 |
|  | ΔANGLE | 0 | 30.85 | 0 | 2.18 | 0 |
|  | ΔDIHED | 0 | 15.41 | 0 | 1.09 | 0 |
|  | ΔUB | 0 | 5.05 | 0 | 0.36 | 0 |
|  | ΔIMP | 0 | 5.35 | 0 | 0.38 | 0 |
|  | ΔCMAP | 0 | 11.8 | 0 | 0.83 | 0 |
|  | ΔVDWAALS | -311.7 | 8.15 | 12.17 | 0.57 | 0.86 |
|  | ΔEEL | 40.67 | 68.42 | 25.05 | 4.83 | 1.77 |
|  | Δ1-4 VDW | 0 | 12.85 | 0 | 0.91 | 0 |
|  | Δ1-4 EEL | 0 | 53.5 | 0 | 3.77 | 0 |
|  | ΔEPB | 120.78 | 42.14 | 25.85 | 2.97 | 1.82 |
|  | ΔENPOLAR | -35.48 | 0.53 | 1.24 | 0.04 | 0.09 |
|  | ΔGGAS | -271.02 | 70.29 | 28.06 | 4.96 | 1.98 |
|  | ΔGSOLV | 85.3 | 42.15 | 25.34 | 2.97 | 1.79 |
|  | **ΔTOTAL** | **-185.73** | **81.96** | **13.73** | **5.78** | **0.97** |

*(a)* The standard deviation (SD) is acquired through the propagation of the uncertainty formula. *(b)* SD represents the sample standard deviation. *(c)* The standard error of the mean (SEM) is determined through the propagation of the uncertainty formula. *(d)* SEM denotes the sample standard error of the mean.

**Supplementary Table S3.** Intermolecular interactions observed in stable and equilibrated AAMD simulation trajectories, analyzed using Arpeggio. The table presents various interaction types, including hydrogen bonding, hydrophobic interactions, polar contacts, van der Waals interactions, proximal interactions, ionic interactions, aromatic interactions, and carbonyl interactions across the tau-tau, FG-Nup98-tau, and FG-Nup98-FG-Nup98 complexes.

| **Types of interaction** | **Tau-Tau**  **complex** | **FG-Nup98-Tau**  **complex** | **FG-Nup98-FG-Nup98**  **complex** |
| --- | --- | --- | --- |
| Hydrogen bonds | 35 | 22 | 19 |
| Weak hydrogen bonds | 20 | 11 | 15 |
| Hydrophobic contacts | 39 | 58 | 80 |
| Polar contacts | 42 | 37 | 27 |
| Weak polar contacts | 35 | 26 | 22 |
| VdW interactions | 21 | 20 | 18 |
| VdW clash interactions | 33 | 25 | 13 |
| Proximal interactions | 1965 | 1719 | 1351 |
| Ionic interactions | 10 | 4 | 0 |
| Aromatic contacts | 0 | 0 | 14 |
| Carbonyl interactions | 5 | 5 | 2 |
| **Total number of interactions** | **2205** | **1927** | **1561** |

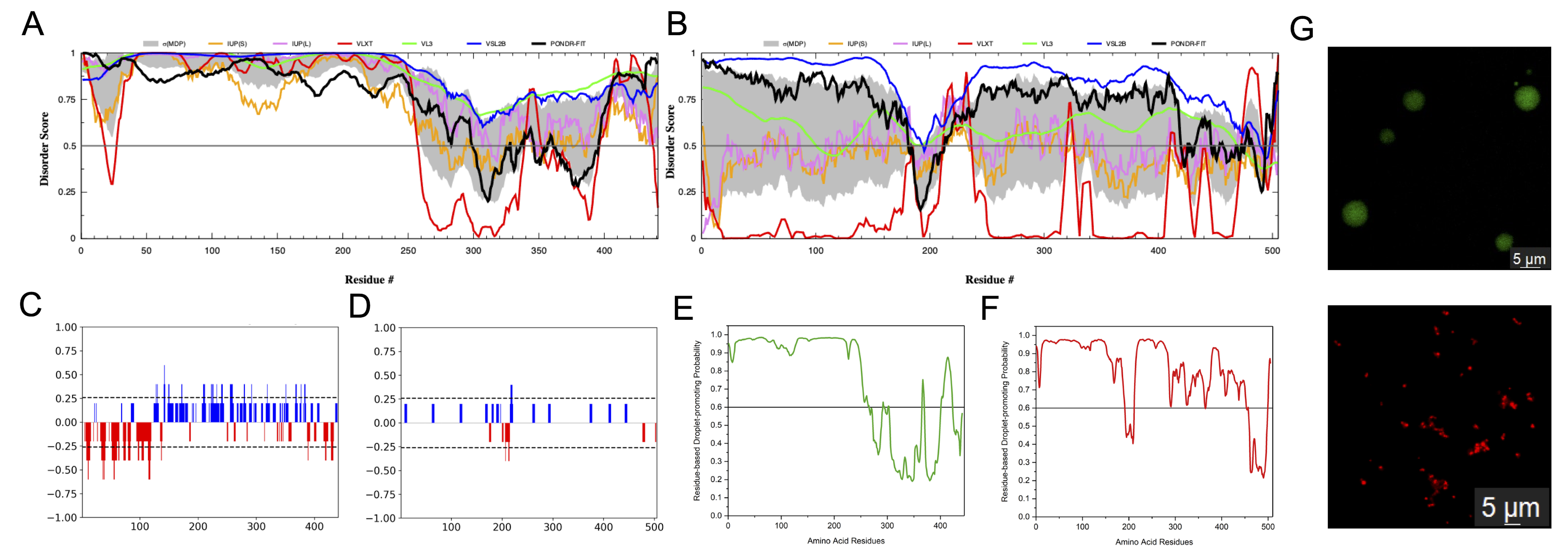

**Supplementary Figure S1: Bioinformatics analysis and phase separation of tau and FG-Nup98.** (A-B) Disorder prediction of tau (A) and FG-Nup98 (B) using AIUPRED. (C-D) Net charge per residue (NCPR) analysis of tau (C) and FG-Nup98 (D) using CIDER. (E-F) Phase separation propensity scores for tau (E) and FG-Nup98 (F) using FuzDrop. (G) Confocal microscopy images confirming phase separation of tau (green) and FG-Nup98 (red) individually.

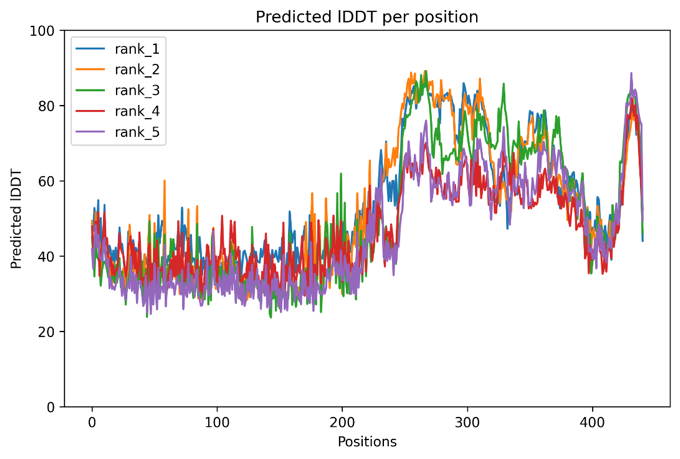

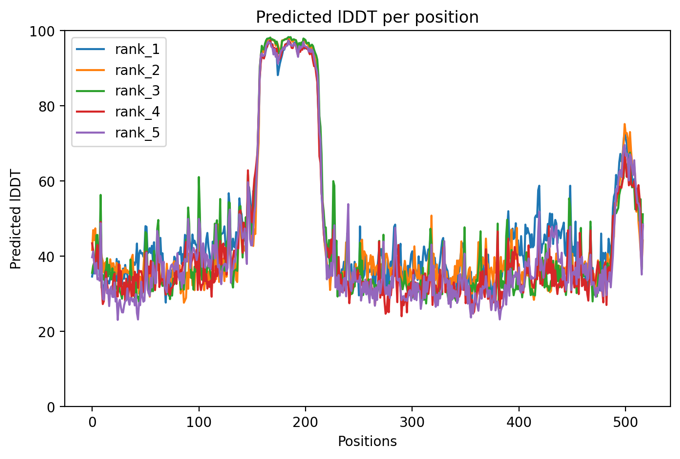

A

B

**Supplementary Figure S2. Predicted Local Distance Difference Test (pLDDT) scores of protein structures generated using AlphaFold v2.0.** (A) pLDDT score distribution for N-terminal tau monomers. (B) pLDDT score distribution for the FG-Nup98 monomer. These scores provide an assessment of model confidence and structural quality.

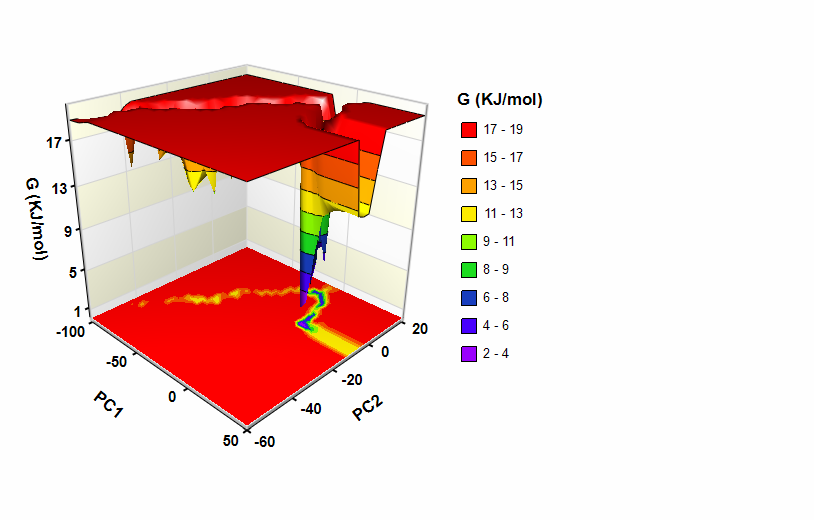

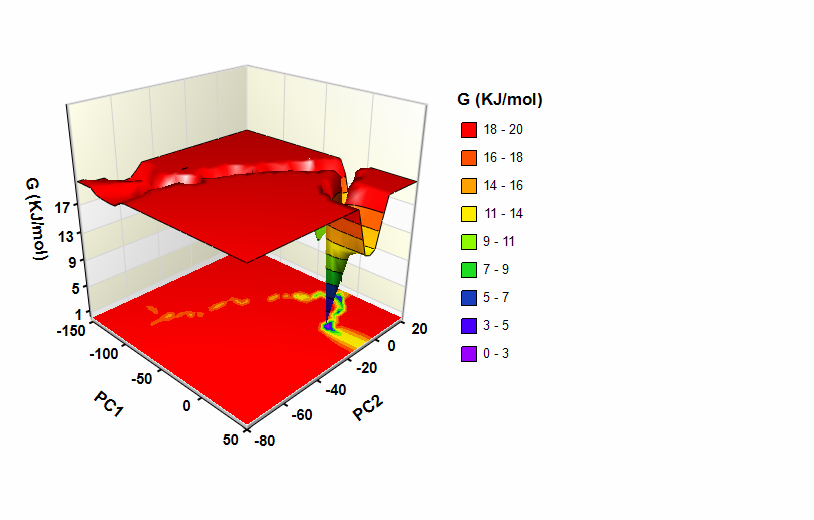

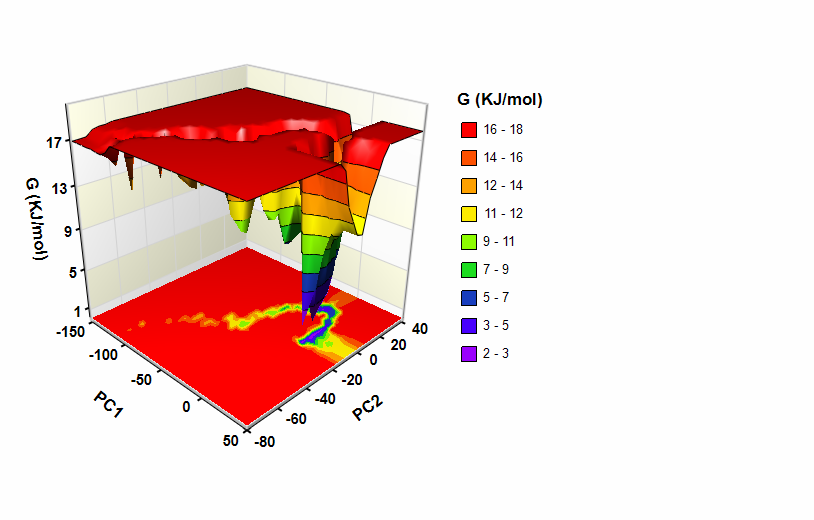

**(D)**

**(E)**

**(F)**

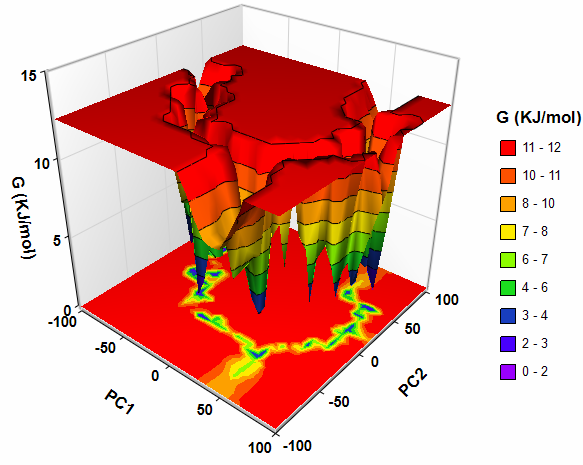

**(A)**

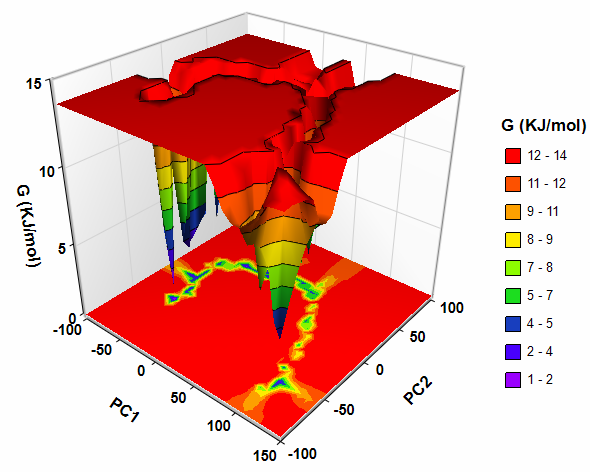

**(B)**

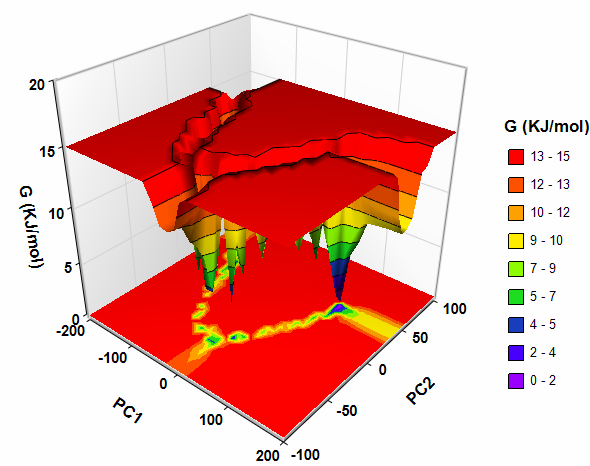

**(C)**

**Supplementary Figure S3. The Gibbs free energy landscapes (FEL) of FG-Nup98-FG-Nup98, FG-Nup98-tau, and tau-tau models derived from AAMD and CGMD simulations.** Panel (A-B-C) depicts the FEL contour plots for FG-Nup98-FG-Nup98, FG-Nup98-tau, and tau-tau models depict the energy distribution. Panel (D-E-F) illustrates the FEL contour plots for the same systems, FG-Nup98-FG-Nup98, FG-Nup98-tau, and tau-tau complexes. The FELs are obtained by projecting the principal components (PC1 and PC2) illustrating the conformational sampling and energy minima. In all FEL plots, the color scheme reflects the energy states: red represents high-energy states, yellow and green indicate low-energy states, while blue and purple correspond to the most stable states with the lowest energy levels. The PCA trace values, measured in nm², reflect the conformational flexibility of the complexes, where lower values indicate greater structural stability. In all-atom molecular dynamics (AAMD) simulations, the trace values are: FG-Nup98-FG-Nup98: **7924 nm²**, FG-Nup98-tau: **7318** **nm²**, and tau-tau: **12163** **nm²**. In coarse-grained molecular dynamics (CGMD) simulations, the values are: FG-Nup98-FG-Nup98: **576 nm²**, FG-Nup98-tau: **905 nm²**, and tau-tau: **1270 nm²**. These findings suggest that the **FG-Nup98-FG-Nup98 and FG-Nup97-tau** complexes exhibit the highest stability, while **tau-tau** is the most flexible and least stable in both simulation models.

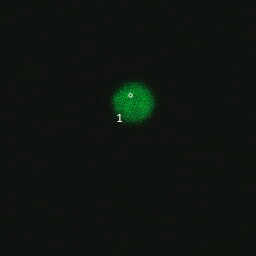

**Supplementary Video 1: FRAP of tau in a colocalized droplet of FG-Nup98 and tau.** The white circle marks the region bleached.

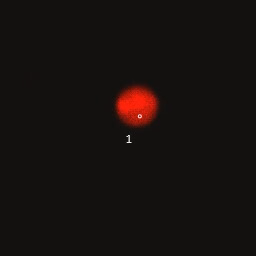

**Supplementary Video 2: FRAP of FG-Nup98 in a colocalized droplet of FG-Nup98 and tau.** The white circle marks the region bleached.

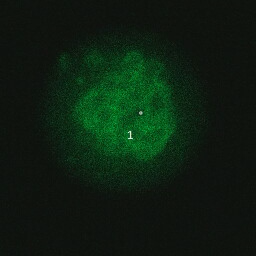

**Supplementary Video 3: FRAP of tau in a nested droplet of FG-Nup98 and tau.** The white circle marks the region bleached.
